# mTORC1 Restricts Hepatitis C Virus Replication Through ULK1-mediated Suppression of miR-122 and Facilitates Post-replication Events

**DOI:** 10.1101/446757

**Authors:** Manish Kumar Johri, Hiren Vasantrai Lashkari, Dhiviya Vedagiri, Divya Gupta, Krishnan Harinivas Harshan

**Affiliations:** CSIR-Centre for Cellular and Molecular Biology, Uppal Road, Hyderabad-500007, India; Academy of Scientific and Innovative Research (AcSIR), Ghaziabad-201002, India

**Keywords:** Hepatitis C Virus, mTOR, Replication, ULK1, miR-122, Autophagy

## Abstract

Mechanistic target of rapamycin (mTOR) is an important kinase that assimilates several upstream signals including viral infection and facilitates appropriate response by the cell through two unique complexes mTORC1 and mTORC2. Here, we demonstrate that mTORC1 is activated early during HCV infection as antiviral response. Pharmacological inhibition of mTORC1 promoted HCV replication as suggested by elevated levels of HCV (+) and (-) RNA strands. This was accompanied by significant drop in extracellular HCV RNA levels indicating defective post-replication stages. The increase in viral RNA levels failed to augment intracellular infectious virion levels, suggesting that mTORC1 inhibition is detrimental to post-replication steps. Lower infectivity of the supernatant confirmed this observation. Depletion of Raptor and ULK1 accurately reproduced these results suggesting that mTORC1 imparted these effects on HCV through mTORC1-ULK1 arm. Interestingly, ULK1 depletion resulted in increased levels of miR-122, a critical host factor for HCV replication, thus revealing a new mechanism of regulation by ULK1. The binary effect of mTORC1 on HCV replication and egress suggests that mTORC1-ULK1 could be critical in replication: egress balance. Interestingly we discover that ULK1 depletion did not interfere with autophagy in Huh7.5 cells and hence the effects on HCV replication and post-replication events are not resultant of involvement of autophagy. Our studies demonstrate an overall ULK1 mediated anti-HCV function of mTORC1 and identifies an ULK1-independent autophagy that allows HCV replication in spite of mTORC1 activation.

## INTRODUCTION

Hepatitis C Virus (HCV) is the major cause of hepatocellular carcinoma worldwide (Choo et al., 1989). It is the lone member of *Hepacivirus* genus under *Flaviviridae* family and is represented by seven genotypes. HCV spreads through blood or organs and over 170 million people are estimated to have been infected by this pathogen (Hadigan and Kottilil, 2011), of which over 71 million people worldwide are chronically infected. Conventional combination therapy using pegylated IFN-α and Ribavirin is associated with inconsistent results and causes severe side effects (Fried, 2002; Manns et al., 2006). In the absence of effective vaccines, recently approved direct acting antivirals (DAAs) have shown remarkable success in virus clearance. However, recent studies indicate the emergence of resistant clones against DAAs (Chiara Di Maio et al., 2017; Takeda et al., 2017). Therefore, there is the need for continued efforts for evolving therapeutic regimens to counter the HCV menace.

Of the several host factors critical for HCV sustenance, miR-122, a liver enriched miRNA, is indispensable for HCV replication (Jopling et al., 2005). miR-122 is a unique case of miRNA mediated positive regulation, where it binds to the 5’ UTR of HCV and stabilizes it (Shimakami et al., 2012). More recent studies suggest that miR-122 does more than providing stability, but is involved in HCV RNA synthesis (Li et al., 2013). Other studies also indicate that miR-122 regulates HCV replication: translation balance (Masaki et al., 2015). It has been demonstrated that simple expression of miR-122 renders otherwise non-permissive hepatic cell lines HCV permissive (Kambara et al., 2012).

Mechanistic target of Rapamycin (mTOR), a major serine/threonine kinase found in eukaryotes is the target of the versatile drug Rapamycin (Brown et al., 1994; Sabatini et al., 1994). mTOR functions as the catalytic subunit of at least two distinct complexes, mTORC1 and mTORC2 (Saxton and Sabatini, 2017). mTORC1 is constituted by Raptor, mLST8, PRAS40 and DEPTOR in addition to mTOR (Kim et al., 2002; Wullschleger et al., 2006). mTORC1 is a major signaling hub that regulates several cellular and metabolic activities such as protein translation, autophagy, lipid and nucleic acid biosynthesis. Deregulated mTORC1 signaling is implicated in several cancers (Hsieh et al., 2010; Laplante and Sabatini, 2012). mTORC1 kinase regulates cell functioning by phosphorylating its diverse set of substrates. Of these, eukaryotic translation initiation factor E (eIF4E)-binding protein (4EBP1) is a major substrate that regulates translation initiation (Haghighat et al., 1995). 4EBP1, in its hypophosphorylated form sequesters eukaryotic cap binding protein eIF4E thereby inhibiting cap dependent translation (Richter and Sonenberg, 2005). However, upon phosphorylation by mTORC1, hyperphosphorylated 4EBP1 loses its affinity for eIF4E thereby facilitating translation initiation. Another well-known substrate of mTORC1 is Unc-51 like autophagy activating kinase 1(ULK1) that activates autophagy. Phosphorylation of ULK1 by mTORC1 at S758 residue inhibits autophagy (Egan et al., 2011). Therefore, active mTORC1 can activate translation initiation and suppress autophagy. However, autophagy is a critical requirement for HCV replication (Dreux et al., 2009; Tanida et al., 2009). Activation of mTORC1 could thus be considered as a potential target for HCV replication by limiting autophagy.

Whereas a wealth of information exists on mTORC1, reports on the regulation by mTORC2 are far fewer. mTORC2 is constituted by its catalytic subunit mTOR, the Rapamycin-insensitive protein of mTOR (Rictor), mLST8, DEPTOR and mammalian stress-activated MAP kinase-interacting protein 1 (SIN1) (Laplante and Sabatini, 2012). mTORC2 is a tyrosine kinase complex (Sarbassov et al., 2005) that controls cell survival and cellular proliferation by phosphorylating several members of the AGC (PKA/PKG/PKC) family of protein kinases (Saxton and Sabatini, 2017). Rapamycin is known to inhibit mTORC1 exclusively during short-term treatment but can inhibit both complexes C1 and C2 during long-term treatment (Sarbassov et al., 2006). Rapamycin has differential effects on mTORC1 substrates and several cancer lines demonstrate resistance to the drug (Choo et al., 2008). Other important active site inhibitors such as Torin1 (Thoreen et al., 2009) and PP242 (Feldman et al., 2009) inhibit both complexes. The latter drugs potently inhibit protein translation even in Rapamycin resistant systems.

mTOR, being a crucial molecule for both translation and autophagy, should be relevant in HCV infection. HCV translation is cap-independent and mediated through internal ribosome entry site (IRES) located in the 5’ UTR of its genome (Tsukiyama-Kohara et al., 1992; Wang et al., 1994). Since cap-independent translation is not directly regulated by mTORC1, changes in mTOR could affect host translation, sparing that of HCV. On the other hand, autophagy is critical for HCV replication and establishment of infection. Regardless of its importance, the role of mTOR in HCV infection has been controversial with certain reports suggesting its essentiality in HCV replication (Stohr et al., 2016) while others suggest its antiviral role (Mannova and Beretta, 2005; Shao et al., 2010; Shrivastava et al., 2012). Several factors including cell types, HCV genotype, MOI, experimental set-up (infection vs. transfection) and duration of infection could have contributed to these divergent conclusions. Our previous studies have demonstrated that HCV protein NS5A activated mTORC1 (George et al., 2012). Notwithstanding the differences, it is evident that mTOR is regulated during HCV infection.

Previously, we had demonstrated that HCV NS5A activated mTOR and cap complex assembly (George et al., 2012) suggesting that mTOR is activated during HCV infection. In the present study, we sought to understand the role of mTOR in HCV life cycle and identified that activation of mTORC1 by HCV early during infection is an antiviral response by the cells. We identify that mTORC1 restricts HCV replication through ULK1 that further executes the regulation through modulating the levels of miR-122. mTOR inhibition reduced viral titer in the supernatant through slower assembly and packaging of the virions. Our studies demonstrate that mTORC1 is key to the balance between viral replication, virion packaging and release. We also identify a disjunction between ULK1 and autophagy in Huh 7.5 cells thereby explaining simultaneous activation of mTORC1 and autophagy in HCV infected cells. Our results indicate a possible molecular explanation of Huh7.5 being the most efficient system for HCV replication.

## Results

### HCV infection activates mTOR

To investigate the regulation of mTOR upon HCV infection, the kinetics of mTOR activity during HCV infection was studied. Huh7.5 cells infected with 0.5 MOI of HCV gt2a were collected at every 24 hr intervals up to 96 hrs (Figure 1A). HCV infection was confirmed by immunobloting of viral core protein. mTOR activity was evaluated by measuring phosphorylation of its substrate 4EBP1. Figure 1B and 1C demonstrated that HCV infection caused remarkable mTOR activation at 24 hpi, as compared with the mock, and moderated later on. Since enhanced mTOR activity was detected early during infection, a probable role of mTOR on HCV was further explored.

**Fig. 1.**
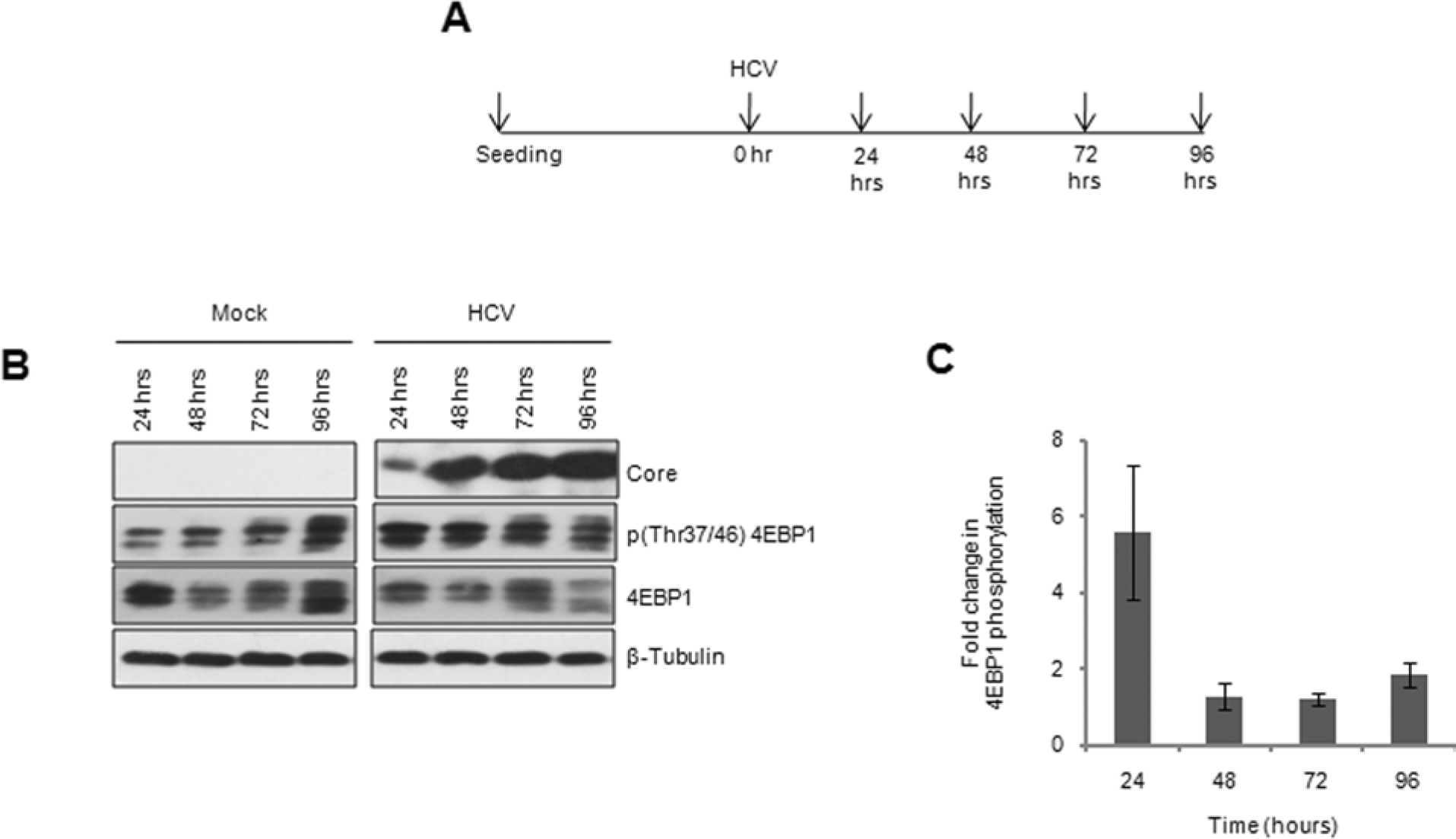
Kinetics of HCV replication and its effect on mTOR activation. (A) Schematic of HCV infection setup. Huh7.5 cells were infected with HCV at 0.5 MOI for 4 hrs after which the infected cells were grown and harvested at every 24 hrs interval. (B) Activation of mTOR during HCV infection in Huh7.5 cells demonstrated by immunobloting of the infected and mock lysates. (C) Quantification of 4EBP1 phosphorylation in HCV infected cells. Densitometric values of phosphorylated 4EBP1 bands from HCV infected samples were normalized to that of the total protein and was further normalized to that of β-Tubulin. These values were normalized to that of the mock-infected samples and fold changes were plotted in graph.

### mTOR restricts intracellular HCV RNA abundance but augments its extracellular levels

Activation of mTOR in response to HCV infection could possibly be either (i) a host antiviral response to HCV replication; or (ii) facilitating HCV infection. We sought to address these possibilities by inhibiting mTOR using two pharmacological inhibitors of mTOR, Rapamycin and Torin1. While mTOR inhibition restrains host translation, it is expected to spare IRES mediated cap-independent HCV translation using. Huh7.5 cells infected with HCV were inhibited with 25nM Rapamycin/750nM Torin1 for 24 hrs starting at 48 hpi (Figure 2A). Prolonged inhibition was important from a clinical management aspect. Both Rapamycin and Torin1, the latter more strongly, inhibited mTOR in HCV infected cells as indicated by dephosphorylations of 4EBP1 and ULK1 (Figure 2B). Inhibitions in mock-infected cells were also observed (data not shown, as they are irrelevant to the study). We studied their effect on HCV by quantifying intracellular HCV (+) and (-) strand abundance. Completion of HCV replication cycle requires generation of both (+) and (-) strands and hence any consistent change in the levels of both strands is a true reflection of changes in replication rate. mTORC1 inhibition is well known to affect protein homeostasis and stability (Zhao et al., 2015). We chose not to use luciferase based replicon assays due to the possibility that mTORC1 inhibitors might affect the stability and enzyme activity of luciferase.

**Fig. 2.**
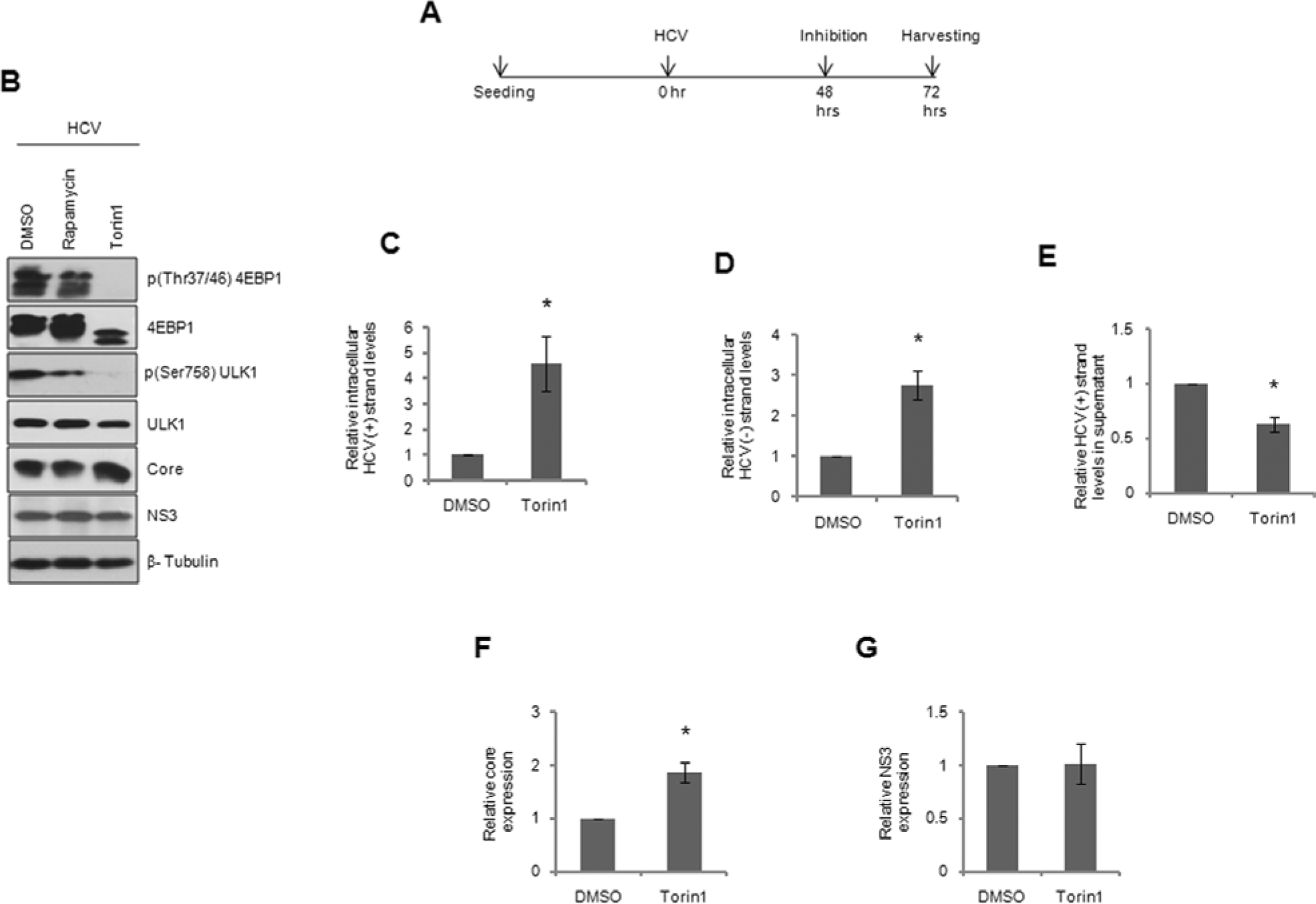
Effect of mTOR inhibition on HCV infection. (A) Schematic depiction of HCV infection and mTOR inhibition. HCV infected cells were inhibited with either Rapamycin or Torin1 at 48 hpi for 24 hrs, after which the cells were harvested. (B) Immunoblot confirming mTOR inhibition by the two inhibitors. Relative intracellular abundance of (C) (+) and (D) (-) strands of HCV, quantified by qRT-PCR. (E) Extracellular HCV RNA abundance in supernatants of inhibited cells. (F) Relative levels of core and (G) NS3 upon mTOR inhibition from immunoblots.

Extensive Torin1 treatment potently induced the levels of HCV (+) and (-) strands (Figure 2C and 2D) and indicated activation of HCV replication. Rapamycin inhibition also caused similar changes in HCV replication (Figure S1A and S1B). Extracellular viral RNA abundance is a measure of HCV particles in the supernatant. Supernatant HCV RNA levels were substantially dropped by Torin1 treatment suggesting that inhibition of mTOR compromises HCV post-replication step such as packaging or egress (Figure 2E). Torin1 treatment did not suppress core or NS3 expression suggesting that HCV replication enhancement did not compromise HCV translation, i.e. not a case of replication-translation switching. (Figure 2B, 2F and 2G). These results indicate that mTOR is an important player in balancing intracellular and extracellular HCV RNA levels.

### Low mTOR activity at the time of infection enhances HCV replication

In previous experiments, the role of mTOR was addressed well beyond the peak HCV replication stage. Since mTOR activation occurred during early hours of infection, we sought to learn the effect of mTOR inhibition at the time of infection on HCV. Pre-inhibition mode is also crucial to understand the effect of mTOR inhibition on HCV entry and replication in previously uninfected liver cells when mTOR inhibitors are part of anti-HCV regimen. Huh7.5 cells were pre-inhibited with Torin1 or Rapamycin for 4 hrs and infected with HCV for 4 hrs in presence of the inhibitors (Figure 3A). Viral supernatants with inhibitors were replaced with growth medium and the cells were allowed to grow further for 68 hrs at the end of which viral RNA levels were assayed. From now on, we refer to this as pre-inhibition mode of infection. Inhibitions at the time of infection and after 4 hrs of infection were confirmed in parallel mock-infected sets (Figure S2A and S2B respectively). Interestingly, mTOR remained inhibited even after 68hrs of treatment (Figure 3B), suggesting that mTOR was constantly under suppression throughout the time duration of infection. Similar to 24 hrs inhibition, pre-inhibition of mTOR by Torin1 substantially increased HCV RNA levels (Figure 3C and 3D) and caused severe reduction in HCV supernatant RNA (Figure 3E), confirming that prolonged suppression of mTOR boosts HCV replication and impairs viral packaging/egress. As in the earlier case, no remarkable drop in core or NS3 expression was observed during pre-inhibition experiments (Figure 3F and 3G), confirming that HCV translation is unaffected by mTOR inhibition. Higher levels of core detected in prolonged-inhibition could be resultant of dynamics of protein stabilization as this was not obvious in pre-inhibition. Rapamycin treatment caused similar effects on HCV replication, albeit less remarkably (Figure S3A and S3B).

**Fig. 3.**
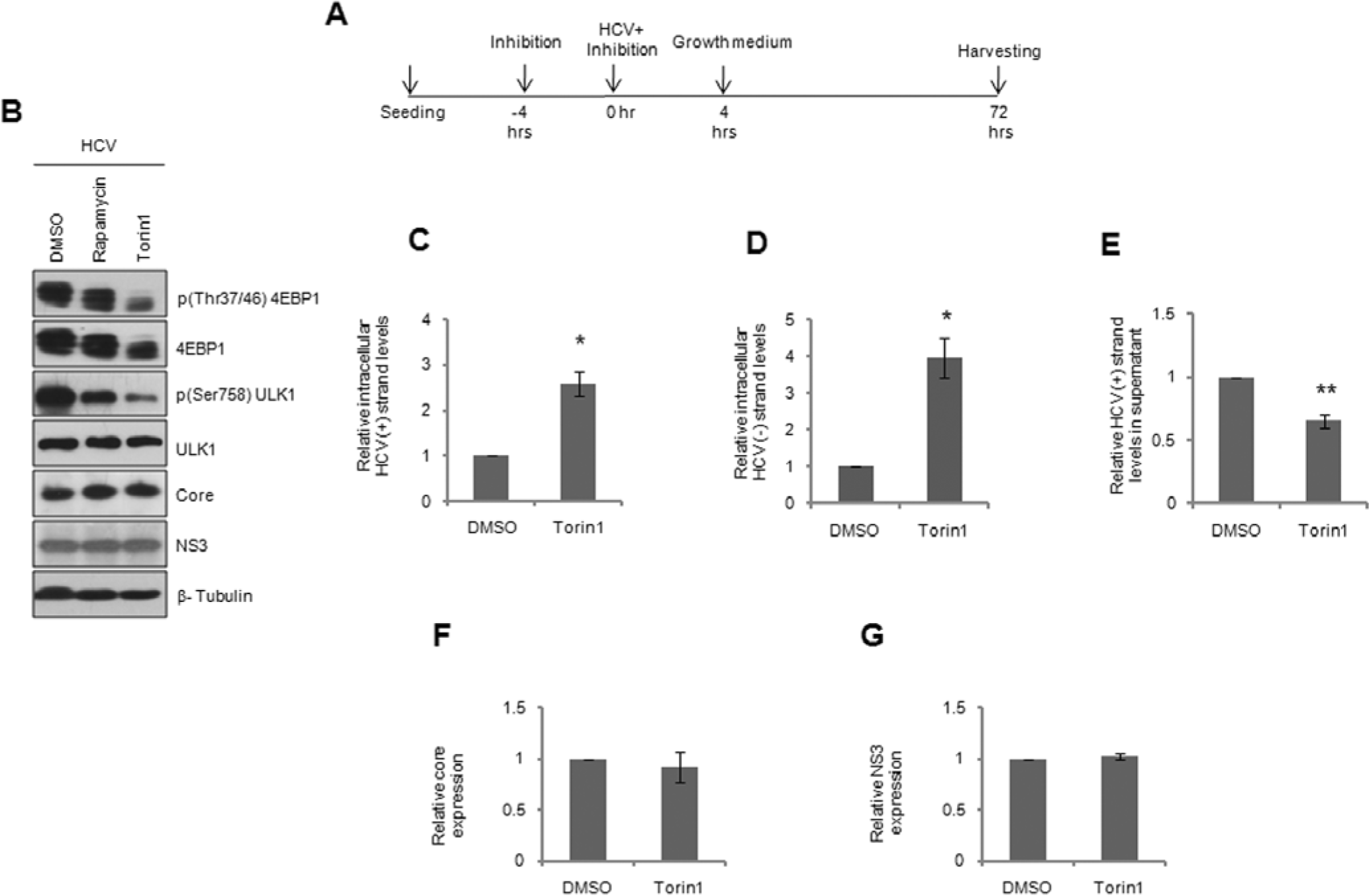
Consequence of pre-inhibition of mTOR on HCV infection. (A) Schematic of mTOR inhibition and HCV infection. Huh7.5 cells were pre-treated with either Rapamycin or Torin1 for 4 hrs prior to HCV infection. At the end of this, cells were infected with HCV in presence of the inhibitors. Subsequently, cells were cultured in fresh media for another 68 hrs at which time they were harvested. (B) Immunoblot validating the inhibition of mTOR. (C) Relative intracellular abundance of HCV (+) and (D) (-) strands, quantified by qRT-PCR. (E) Extracellular HCV RNA levels quantified in supernatants. (F) Relative levels of core and (G) NS3 upon mTOR inhibition from immunoblots.

### mTOR inhibition suppresses HCV assembly and viral release

To address whether mTOR inhibition affects the viral post-replication steps leading to loss in infectivity, intracellular viral particles extracted from equal number of pre-treated, HCV infected cells used in experiments shown in Figure 3 and their infectivity was tested on naïve Huh7.5 cells. Infectivity between the samples was measured by comparing the HCV RNA levels in the infected cells and their differences were represented as relative infectivity. Notwithstanding the higher RNA presence, infectivity of cellular virions extracted from Torin1 treated cells did not vary from that from DMSO treated samples (Figure 4A). The results suggest that despite augmenting intracellular RNA abundance, Torin1 treatment did not alter the infectious viral particle number. Thus, mTOR inhibition promotes HCV replication but not viral assembly. Next, we tested the infectivity of equal volumes viral supernatants from Torin1 treated cells by infecting naïve cells. This was again measured by relative HCV RNA levels in the infected cells. Interestingly, infectivity of HCV supernatant from Torin1 treated cells was substantially lower as compared with DMSO treated samples (Figure 4B). The drop in infectivity was well comparable with that of HCV RNA levels in supernatants following Torin1 treatment (Figure 3E). Comparable results were observed in prolonged treatment as well (Figure S4A and S4B). This result suggests that the virions released in Torin1 treated cells are indeed infectious, but their release into the supernatant is defective. This presented a conundrum because our earlier results suggested that viral assembly seemed unaffected (Figure 4A). However, the drop in supernatant viral particles is a clear indicator of reduced assembly or release. Any inhibition of release, but not assembly, would have resulted in intracellular accumulation of infectious virions leading to increased infectivity by intracellular virions that was not the case (Figure 4A). HCV assembly, packaging and release are integrally associated and our results suggest that mTOR inhibition activated HCV replication and suppressed later events of packaging and release. Further focused studies are required to delineate these processes.

**Fig. 4.**
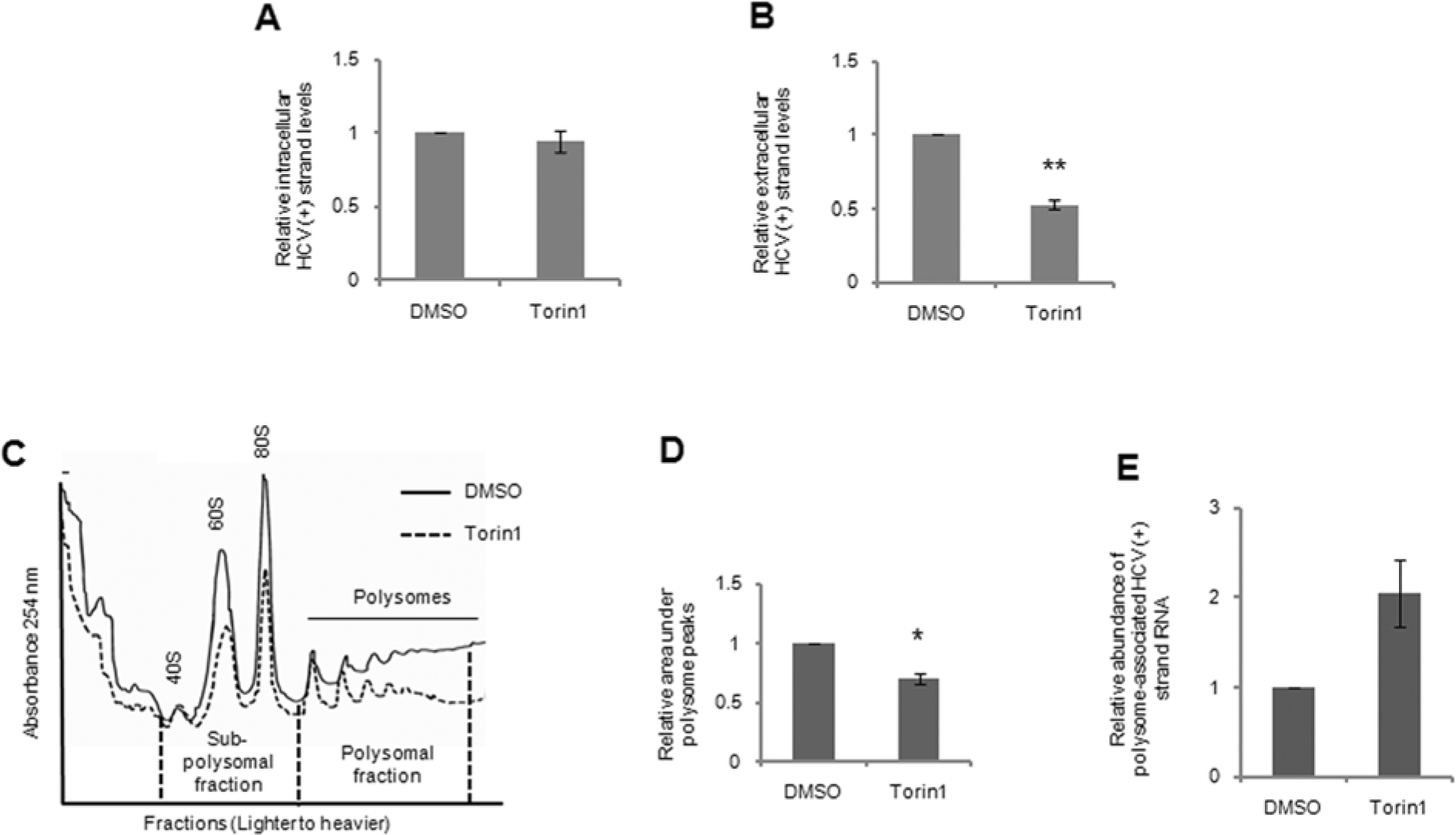
HCV post-replication steps are perturbed with Torin1, but not viral protein translation. (A) Equal number of HCV infected Huh7.5 cells (primary infection) in the presence of Torin1 or DMSO from Figure 3A were freeze-thawed and the released viral particles were resuspended in equal volumes of serum-free media. This viral suspension was used to infect fresh Huh7.5 cells for 72 hrs (secondary infection) after which intracellular HCV RNA abundance was quantified. Relative HCV (+) strand abundance from Huh7.5 cells infected in the second round, infected with intracellular virus of either Torin1 or DMSO treated cells during the primary infection. (B) Relative HCV (+) strand abundance from Huh7.5 cells infected with the supernatant of Torin1 or DMSO treated cells during the primary infection as shown in Figure 3E. (C) Polysome profiles of HCV infected cells after pre-treatment with Torin1. Cells harvested at 72 hpi in pre-treatment mode were lysed and polysome profiles were generated. (D) Quantitative representation of relative polysome association in Torin1 and DMSO treated cells as described in (C). Area under the polysomal fractions were measured for representation. (E) All the polysome fractions were pooled, RNA was precipitated and relative association of HCV RNA with polysome fractions of Torin1 pre-treated cells against DMSO treated control cells.

mTORC1 is not known to regulate cap-independent translation. In prolonged treatment cases, Torin1 induced core levels while no change was observed in pre-treatment (Figures 2B, 2F, 3B and 3F). Levels of another viral protein, NS3, were unchanged in both conditions (Figures 2B, 2G, 3B and 3G). These findings suggested that mTORC1 inhibition did not suppress HCV translation. This was verified by measuring HCV RNA loading in polysome during Torin1 treatment. In contrast to the reduction in polysome size (Figure 4C and 4D), HCV RNA association with the polysomes was enhanced (Figure 4E) by Torin1 pre-treatment, confirming the previous reports that mTOR inhibition does not inhibit IRES translation. Therefore, the drop in HCV RNA and infectivity in the supernatant upon Torin1 treatment is not due to perturbation of HCV translation, but possibly due to assembly and packaging.

### mTORC1, but not mTORC2, restricts intracellular HCV RNA abundance

Torin1 inhibits both mTORC1 and mTORC2. Rapamycin inhibits only mTORC1 during short treatment but can inhibit both mTORC1 and mTORC2 upon extensive treatment. The effect of mTOR inhibition on HCV could have been mediated through either of the mTOR complexes. In order to identify the mTOR complex involved in HCV regulation, we depleted Raptor, a major regulatory component of mTORC1, in Huh7.5 cells by siRNA prior to HCV infection (Figure 5A). Raptor depletion resulted in lower 4EBP1 and ULK1 phosphorylations as expected (Figure 5A). Similar to long-term inhibition with Torin1 and Rapamycin, Raptor depletion induced intracellular HCV RNA levels (Figure 5B) and suppressed HCV RNA abundance in the supernatant (Figure 5C). On the contrary, depletion of Rictor, an integral mTORC2 component, had no effect on HCV replication (Figure S5A and S5B). These results confirm that mTOR restricts HCV replication and promotes viral release through mTORC1 but not mTORC2.

**Fig. 5.**
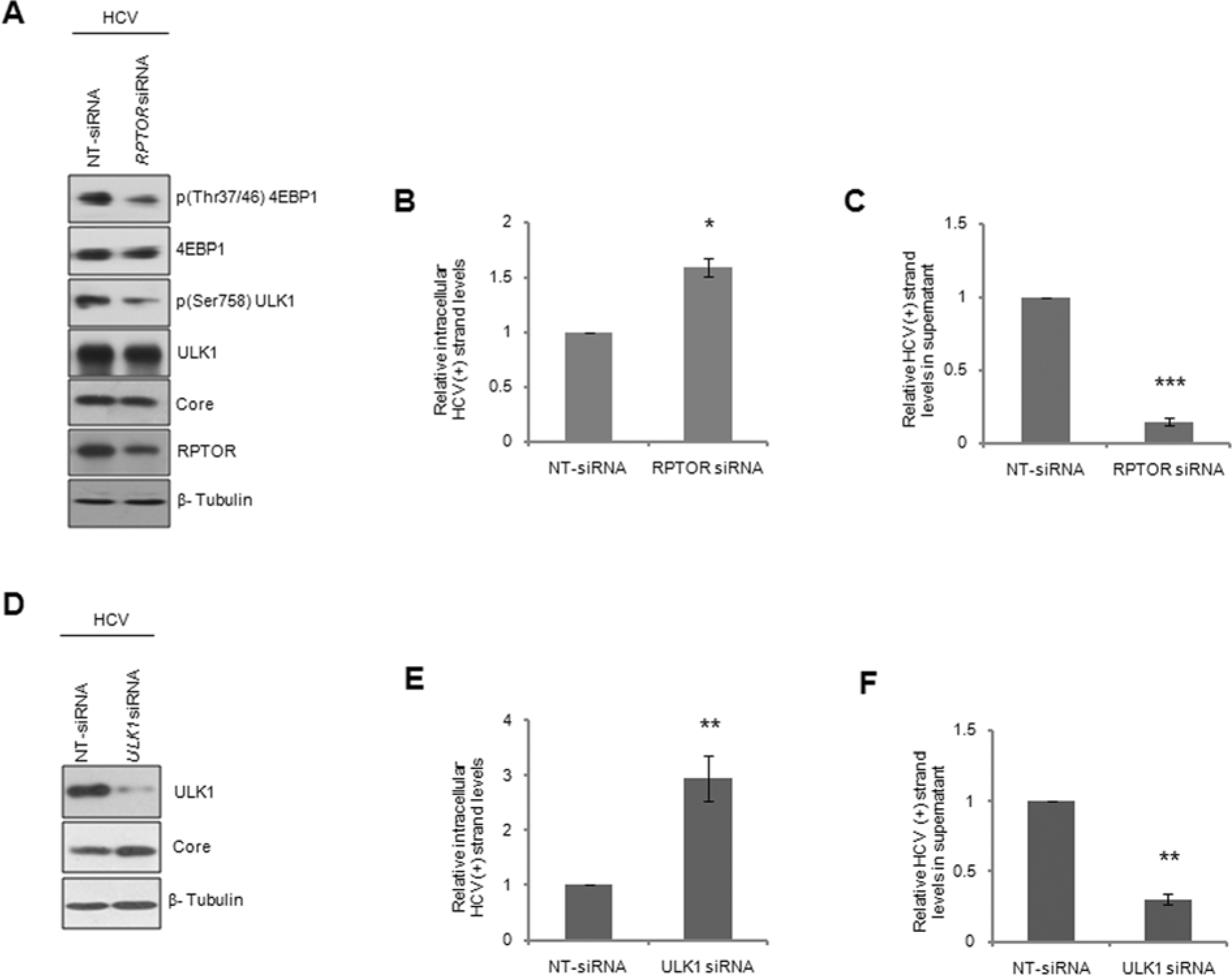
mTORC1 restricts intracellular HCV RNA abundance and facilitates extracellular HCV RNA levels through ULK1. (A) Confirmation of Raptor depletion by immunobloting. Huh7.5 cells were transfected with siRNA pool targeting *RPTOR* or with non-targeting siRNA 18 hrs before infection. 72 hpi, cells were harvested for analysis. (B) Relative intracellular abundance of HCV (+) strands quantified by qRT-PCR. (C) HCV abundance quantified in supernatants. (D) Immunobloting for confirmation of ULK1 depletion. Huh7.5 cells were transfected with siRNA pool targeting *ULK1* or with non-targeting siRNA 18 hrs before infection. 72 hpi, cells were harvested for analysis. (E) Relative intracellular abundance of HCV (+) strands quantified by qRT-PCR. (F) HCV RNA levels quantified in supernatants.

### mTORC1 restricts intracellular HCV RNA levels through ULK1

4EBP1 and ULK1 are key substrates of mTORC1. We sought to identify the mTORC1 substrate through which mTORC1 regulated HCV replication. Depletion of 4EBP1 (Figure S6A and S6B), did not alter HCV replication in Huh7.5 cells, suggesting its dispensability in HCV replication. Next, we analyzed HCV replication in ULK1 depleted Huh7.5 cells. Interestingly, robust ULK1 depletion (Figure 5D) caused significant increase of intracellular (Figure 5E) and severe drop of supernatant HCV RNA levels (Figure 5F), consistent with mTOR inhibition and Raptor depletion. Taken together, these results demonstrate that mTORC1 restricts HCV replication through a mechanism that involves ULK1, but not 4EBP1.

### mTORC1 regulates miR-122 levels through ULK1

Since miR-122 is indispensable for HCV replication, we next enquired its status during mTORC1 inhibitions and knock-downs. Torin1, in both pre-treatment and prolonged treatment resulted in considerable increase in miR-122 levels in HCV infected cells (Figures 6A and S7A respectively) and during ULK1 and Raptor depletions (Figure 6B and S7B respectively). Introduction of anti-miR-122, but not the control anti-miR, efficiently removed miR-122 (Figures 6C and 6D) and caused substantial drop in HCV (+) and (-) strand RNA suggesting that the over-expressed miR-122 was functional (Figures 6E and 6F). HCV translation was suppressed under these conditions as well (Figure 6C). These observations suggest that mTORC1 regulates miR-122 levels through ULK1. These data also suggest that ULK1 has an unidentified role of regulating miR-122 levels through which HCV replication is regulated. These results propose that mTORC1, through ULK1, inhibits miR-122 levels. To the best of our knowledge, this mechanism of regulation of HCV replication is hitherto unidentified.

**Fig. 6.**
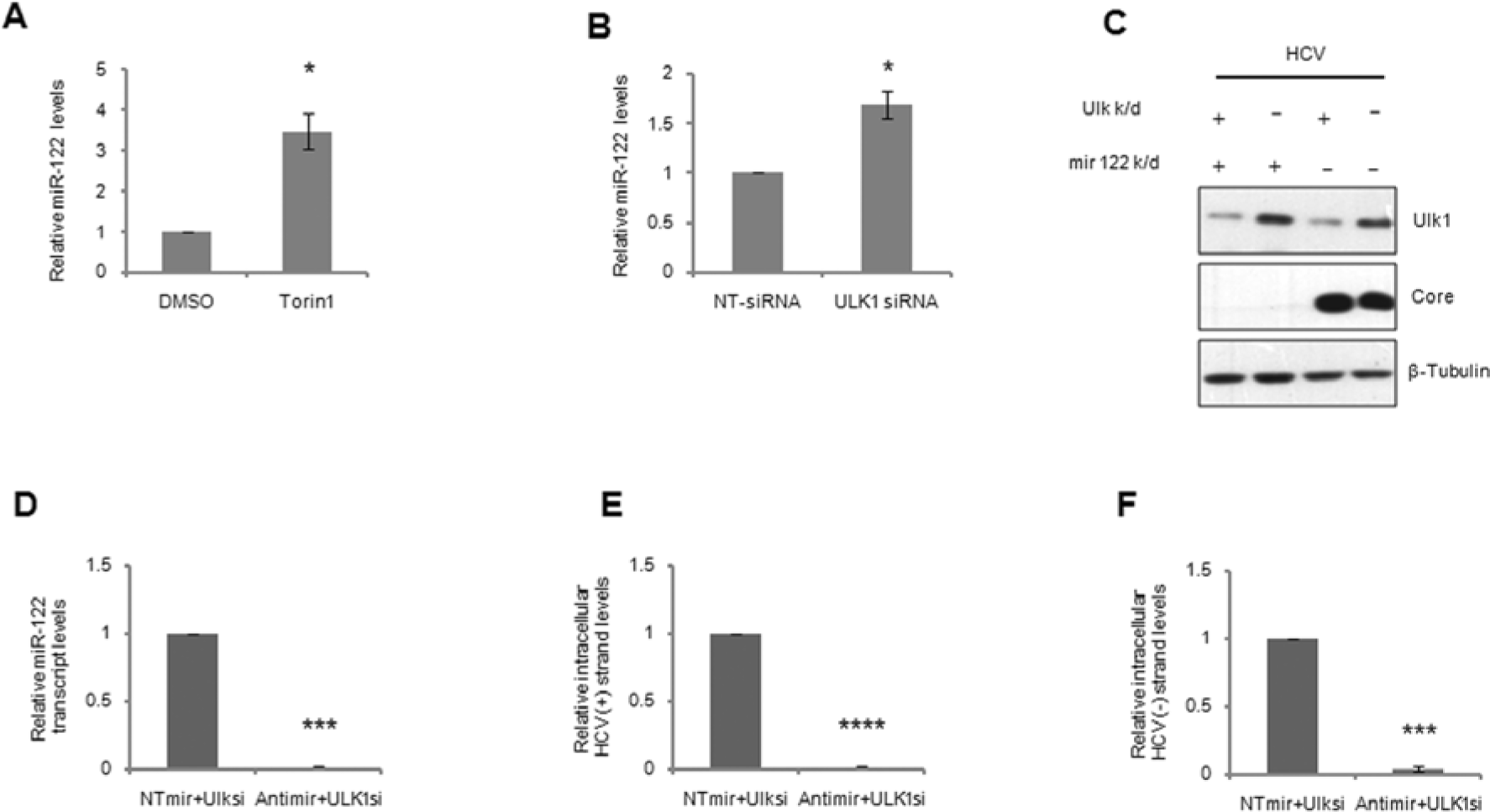
mTOR restricts miR122 transcription through ULK1. Relative miR-122 levels in HCV infected cells subjected to (A) Torin1 pre-treatment and (B) ULK1 depletion. The miR-122 levels were quantified by Taqman based kits and normalized against *RNU6B*. (C) Effect of anti-miR-122 on ULK1 mediated activation of miR-122 in HCV infection. Confirmation of ULK1 knockdown and HCV infection by immunobloting. Huh7.5 cells were transfected with siRNA pool targeting *ULK1* or with non-targeting siRNA 18 hrs before infection. 72 hpi, cells were harvested for analysis. 24 hours prior harvesting 200uM anti-miR122 was transfected. The miR-122 levels were quantified by Taqman based kits and normalized against *RNU6B*. (D) Quantitation of miR-122 levels after transfecting anti-miR-122 or the control anti-miR. (E) Relative intracellular abundance of HCV (+) strands and (F) (-) strands in the presence of antimiR-122.

### Uncoupling of ULK1 with autophagy facilitates maintenance of autophagy even under mTOR activation

Our results have presented a conundrum since mTORC1 activation would normally suppress autophagy thereby causing a probable inhibition of HCV replication. To address this important point, the status of autophagy was analyzed in Torin1 pre-treated samples described in Figure 3A. Pre-treatment with Torin1 of HCV infected cells induced autophagy, evident from the depleted p62 levels (Figure 7A), thus validating the anti-autophagic role of mTOR. Thus, autophagy activation by Torin1 could be a possible contributor to the enhanced HCV replication. However, this analysis was performed at the end of 72 hours of infection and hence is not a reflection of the status of autophagy at the time, and during the early hours of infection that is critical for establishment of HCV infection. To ascertain this, we treated Huh7.5 cells with Torin1 for 8 hrs, as described in Figure S2B, at the end of which cells were analyzed for autophagy. Not surprisingly, activation of autophagy was also evident at the end of 8 hrs of Torin1 treatment without HCV infection (Figure 7B), indicating that this activation could possibly have contributed to the elevated intracellular HCV strands. Similar induction of autophagy was found in prolonged inhibition (Figure 7C). Next, autophagy markers were evaluated in ULK1 depleted cells upon HCV infection. Surprisingly, ULK1 depletion caused no appreciable change in the levels of p62 and Beclin and in LC3 lipidation (Figure 7D-F). To assess the contribution of autophagy in the Torin1 mediated activation of HCV replication, we pre-treated cells with Chloroquine for 6 hrs in presence or absence of Torin1. Treatment was followed by HCV infection for 4 hrs in presence of the inhibitors as indicated above. Chloroquine mediated autophagy inhibition, confirmed by the accumulation of p62 and lipidation of LC3A/B (Figure 7G), caused severe inhibition of HCV replication (Figure 7H). Similar pre-treatment with Bafilomycin also caused substantial drop in HCV RNA abundance (Figure S8), confirming that inhibition of autophagy overrides activation of HCV replication by Torin1. However, since ULK1 depletion enhanced HCV replication in the absence of autophagy induction, we propose that autophagy induction during mTORC1 inhibition could be through ULK1 independent mechanisms. We speculate that mTOR mediated restriction on HCV would have been more effective if autophagy was under regulation by ULK1 in this system. Our results demonstrate that mTORC1 restricts HCV replication through ULK1 mediated mechanism without restricting autophagy and this uncoupling allows HCV to flourish in Huh7.5 cells.

**Fig. 7.**
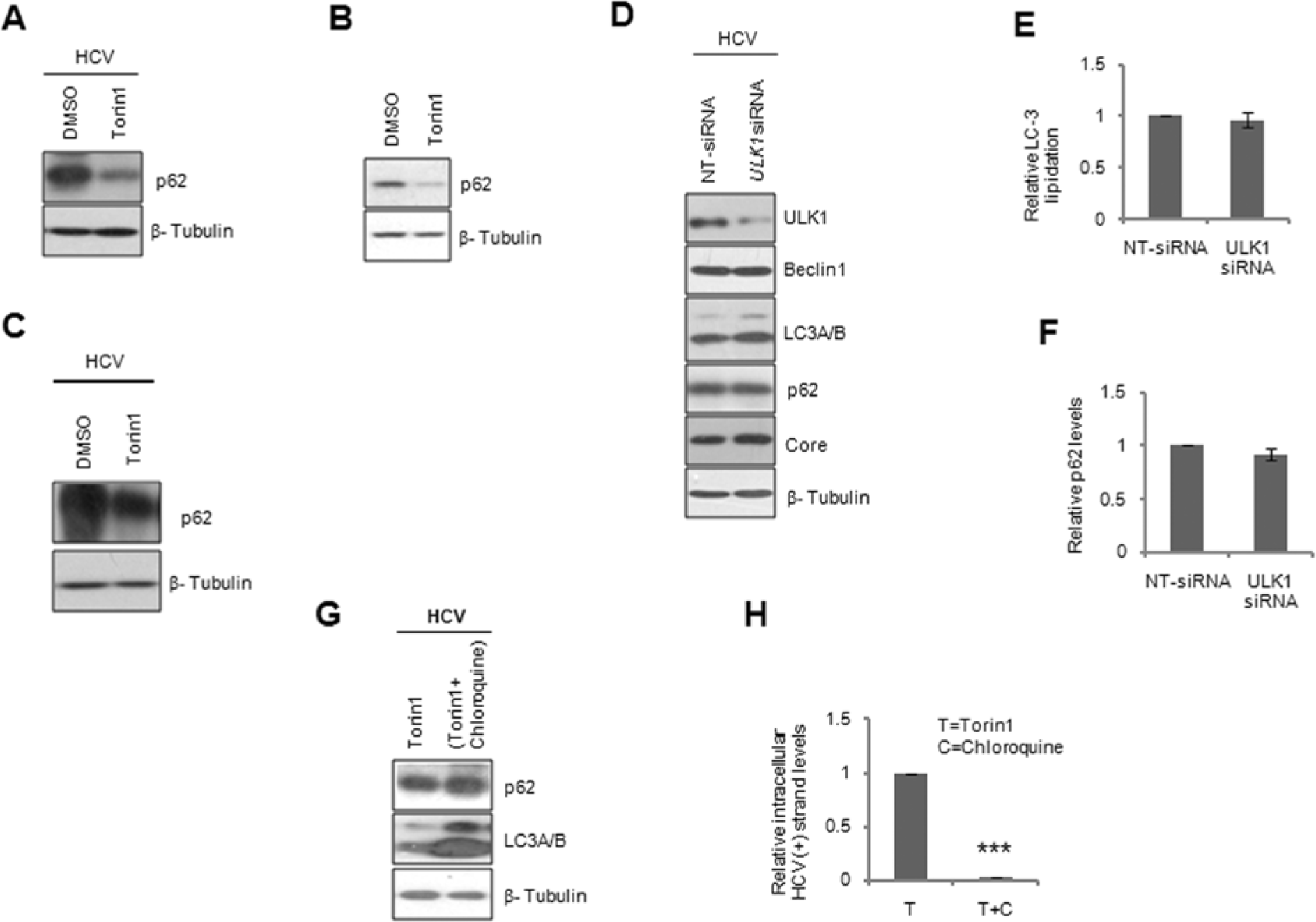
ULK1 does not regulate autophagy in Huh7.5 cells upon HCV infection. (A) Analysis of p62 levels following Torin1 treatment of HCV infected Huh7.5 cells in pre-treatment mode by immunobloting. (B) Evaluation of autophagy activation in Huh7.5 cells treated with Torin1 for 8hrs. Cells treated with Torin1 for 8 hrs were harvested to analyze autophagy markers by immunobloting. (C) HCV infected cells were inhibited with Torin1 as shown in Figure 2. Immunoblot of p62 after 24 hours of Torin1 post-inhibition. (D) Investigation of autophagy in ULK1 depleted Huh7.5 cells upon HCV infection as described in the Figure 5A, by immunobloting. (E) Graphical representation of relative LC3A/B lipidation and (F) that of p62 levels in samples displayed in Figure 7D. (G) Analysis of autophagy in Huh7.5 cells pretreated with Torin1 in presence or absence of Chloroquine, by immunobloting. Huh7.5 cells were pre-treated with 750nM Torin1 with or without 25µM Chloroquine for 6 hrs after which they were both infected with HCV for 4 hrs in the presence of the inhibitors. Post-infection, cells were allowed to grow in fresh media for another 68 hours before harvesting. (H) Graphical representation of relative HCV intracellular (+) strand titers quantified by qRT-PCR. T =Torin1, C=Chloroquine.

## DISCUSSION

The current work identifies two critical functions of mTOR on HCV life cycle. On one hand, it restricts HCV replication and on the other, it facilitates steps involved in the assembly of the virions and their release. The dual effects of mTORC1 on HCV are clearly interpreted from the inhibitor and knockdown studies. mTORC1 activation early during HCV infection could have been interpreted either as a pro-viral, pro-growth response or as an antiviral response. Since mTOR restricts HCV replication that precedes the packaging, we conclude that mTOR is part of anti-HCV machinery in hepatoma cells.

The increase in (+) strands of HCV following long-term inhibition could be resulted by increased replication rate, by accumulation of viral genome/packaged virions or by replication-translation switch. However, the concurrent increase in the (-) strand of HCV confirmed that mTORC1 inhibition indeed activated its replication. Strand-specific q-RT PCR is a powerful technique in determining the replication status of (+) stand RNA viruses through abundance of (-) strand RNA (Komurian-Pradel et al., 2004; Plaskon et al., 2009; Rance et al., 2012).

Effect of mTOR on HCV replication is addressed by previous studies. However, annotating mTOR as a supportive or a suppressive host factor for HCV replication remained debatable. Present study recommends mTOR as an antiviral host factor that limits HCV replication. Our results of HCV mediated suppression of HCV replication corroborate with certain previous report (Mannova and Beretta, 2005) and contradicts with a few others (Huang et al., 2013; Stohr et al., 2016). Several factors including cell types, HCV genotype, MOI, experimental set-up (infection vs. transfection) and duration of infection could have contributed to these divergent conclusions. However, our studies have looked at the role of mTOR comprehensively and used infectious viral system to reach our findings.

Infectivity assay using the intracellular virions of Torin1 treated samples from the first step indicates that post replication steps of HCV cycle remain perturbed by mTOR inhibition. Increase in viral replication due to drop in viral translation (replication-translation switch) is shown in many RNA viruses including HCV (Ray and Das, 2011). However, mTOR inhibitions in our study did not depict any drop in HCV core and NS3 levels suggesting that mTOR inhibition does not trigger a replication: translation switch in favor of replication by suppressing HCV translation. The increase in intracellular HCV RNA abundance upon mTORC1 inhibition is clearly the outcome of increased replication.

Our studies unearthed previously unknown mechanism of regulation of antiviral activities of mTORC1 and identified a new member in this pathway, ULK1. Since ULK1 depletion caused activation of miR-122 transcription, it could be suggested that ULK1 regulates their transcription. It is possible that ULK1 mediates transcriptional regulation through other molecules or directly involved with their transcription. Nuclear functions of ULK1 have been described recently (Joshi et al., 2016).

Egress is a larger process that includes packaging of the genome and its transport through ER-Golgi system before the release takes place. HCV particle assembly is associated with lipid droplets **(Wendel et al.)** in addition, requires factors that are necessary for VLDL assembly and apolipoprotein generation **(Gastaminza et al., 2008; Herker et al., 2010)**. HCV trafficking is dependent on ER-Golgi trafficking system **(Mankouri et al., 2016)** and microtubule polymerization **(Lai et al., 2010)**. Since mTORC1 regulates lipid metabolism **(Laplante and Sabatini, 2009)** and trafficking **(Jiang and Yeung, 2006)**, the assembly and trafficking of HCV particles is a likely target of mTORC1 inhibition. In support, recent reports implicate the roles of ULK1 in lipid metabolism **(Ro et al., 2013)** and cytoskeletal rearrangements **(Caino et al., 2013)**. At this stage, the mechanistic details on the reduced HCV RNA abundance in the supernatant of mTOR inhibited cells are unclear.

mTOR inhibition caused significant cell death in pre-inhibition and prolonged inhibition (Figures S9 A-D). Earlier studies have demonstrated an association between cell division and HCV replication. However, the inhibitory effect of the inhibitors on cell division does not contribute significantly to HCV stages as ULK1 and Raptor knockdown caused similar effects on HCV stages without causing any cell division effects (Figures S9E and S9F.

Induction of mTORC1 is antagonistic to autophagy. ULK1 knockdown has been known to block autophagy in HEK293 and MEFs (Chan et al., 2007)Huh7.5 cells is the only known robust infection system for HCV. A single point mutation in the dsRNA sensor retinoic acid-inducible gene-I (RIG-I) proposed to play a role in higher permissiveness for HCV RNA replication (Sumpter et al., 2005). However, we propose that uncoupling of mTORC1 with autophagy facilitated active replication of HCV in Huh7.5 cells. ULK1 depletion unquestionably had no effect on autophagy markers in Huh7.5 cells. We hypothesize that though mTORC1 phosphorylates ULK1 during the initial hours of infection, it spares autophagy due to the uncoupling of ULK1 with autophagy. We propose that this could be a major factor that allows high permissivity for HCV in Huh7.5 cells. In systems where ULK1 still regulates autophagy, activation of mTOR might effectively suppress autophagy thereby preventing HCV replication. In agreement, Shrivastava et al. identified concurrent activation of autophagy and mTOR (Shrivastava et al., 2012) and concluded that HCV induces autophagy while activation of mTOR facilitated hepatocyte proliferation. Since autophagy seems independent of ULK1 in Huh7.5, HCV is able to maintain its replication even in the presence of activated mTORC1.

### Conclusion

Our studies demonstrate that mTOR is activated during the initial stages of HCV infection. mTOR suppression enhanced HCV RNA titer generated by augmenting replication and hence mTOR activation can be considered as an antiviral response. Simultaneously, mTOR inhibition lowered extracellular HCV levels through suppressing viral package or egress, a feature through which mTOR assumes a virus friendly molecule. mTOR executed these effects through ULK1 and regulated HCV replication through modulating the levels of miR-122. Interestingly, ULK1, though a key molecule in autophagy initiation, does not regulate autophagy in Huh7.5 cells and hence the anti-HCV effects of mTOR could not impart its restriction on HCV replication through inhibiting autophagy. Thus, this uncoupling allows HCV to replicate by utilizing autophagy even in the background of active mTOR.

## Experimental procedures

### Antibodies and inhibitors

All antibodies except p62 and β-Tubulin (Thermo Fischer Scientific) and HCV core (Abcam), were purchased from Cell Signaling Technology. Torin1 was bought from Tocris Bioscience and Rapamycin was from Merck Millipore. Chloroquine and Bafilomycin were from Sigma Aldrich. siRNA pools against RAPTR and ULK1 were from Dharmacon. *4EBP1* and *RICTOR* shRNAs were from Thermo Fisher Scientific.

### Generation of HCVcc particles and titration

Human Hepatoma cell line, Huh7.5 was cultured as described (George et al., 2012). HCV infectious particles were prepared by *in vitro* transcription (IVT) of full-length genome from pFL-J6/JFH1, followed by transfection into Huh7.5 cells (George et al., 2012; Lindenbach et al., 2005). Supernatants from transfected cells were collected, filtered through 0.45 µm filter and quantified by qRT-PCR by absolute quantification. Briefly, RNA was isolated from viral supernatant and reverse transcribed using HCV RT Rev primer (5’-TGCACGGTCTACGAGACCTC-3’). HCV genome was quantified using HCV SG1F (5’-TATGCCCGGCCATTTGGGCG-3’) and HCV SG1R (5’-TACGAGACCTCCCGGGGCAC-3’) primers and the viral RNA abundance was estimated as copies/ml.

### Primary infection and replication assay by qRT-PCR

Huh7.5 cells were infected with 0.5 MOI of HCV for four hours followed by the replacement of the virus with fresh media until the cells were harvested. HCV (+) and (-) RNA strands were quantified by qRT-PCR. Briefly, infected cells were harvested and total RNA was prepared using Nucleospin RNA kit (Macherey-Nagel). Total RNA from the supernatants was prepared using Trizol (Thermo Fisher Scientific). Total RNA from infected cells or supernatants were converted into cDNA using HCV strand-specific primer (5’-TGCACGGTCTACGAGACCTC-3’ for +ve and 5’-ATGAATCACTCCCCTGTGAG-3’ for –ve strand). HCV specific regions were amplified from equal amounts of cDNA using SYBR Premix Ex Taq (Takara Bio Inc.) in Lightcycler 480 (Roche Molecular Diagnostics). Fold changes between samples were calculated by using ΔΔ Cp values with endogenous GAPDH control.

### HCV infectivity assay

Infectivity of HCV particles from the infected cells or cell supernatants was measured by infecting naïve Huh7.5 cells with them. First, Huh7.5 cells were infected with HCV with Torin1 or DMSO treatment for 72 hrs. For measuring the infectivity of intracellular virions, equal numbers of cells from different treatments were freeze-thawed thrice in serum-free DMEM. Equal volumes of these virion containing media were used to infect naïve Huh7.5 cells for 72 hrs. Subsequently, total RNA was prepared from these infected cells and HCV RNA levels were quantified by qRT-PCR as described earlier. Relative HCV RNA levels are reflection of infectivity of virions from the sample and were plotted in graphs. For measuring the infectivity of supernatants generated from the primary cultures subjected to HCV infection and treatment, equal volumes of the supernatants were used to infect naïve Huh7.5 cells. 72 hrs later, total RNA was prepared as described in earlier sections and HCV RNA was quantified by qRT-PCR. Here again, relative HCV RNA levels represented the infectious viral units in the supernatant and thus graphically represented.

### Real-time quantitative RT-PCR (qRT-PCR)

Total RNA prepared from samples were converted to cDNA using Primescript RT and random hexamer. qPCR was performed using SYBR Premix Ex Taq and gene specific primers. *HNF4A* quantification was described elsewhere (Parveen et al., 2017). miRNAs were prepared using Nucleospin miRNA kit (Macherey-Nagel). miR-122 was quantified using Taqman Gene Expression Assay kits from Thermo Fisher Scientific and normalized against *RNU6B*. All the oligos were procured from Bioserve Bio Technologies. Oligo details are provided in Table S1.

### Immunobloting

Immunobloting was performed as described earlier (George et al., 2012). Densitometric analyses of blots were done by using Image J software. Relative phosphorylations were calculated from the ratios of any phosphorylated band to its total protein that was normalized to the corresponding β-Tubulin as loading control (P/T/C).

### Polysome analysis

Polysome profiles of HCV infected cells undergone pre-treatment were prepared as described elsewhere (George et al., 2012). Polysomal fractions from DMSO and Torin1 pre-treated HCV infected samples were processed further to prepare polysomal RNA. Briefly, pooled fractions were incubated with 3M sodium acetate and ethanol and RNA was precipitated. To remove Heparin from the preparation, precipitated RNA was incubated with 2.5M lithium chloride for 2 hrs at -30°C. RNA was precipitated at maximum speed and purified using Nucleospin RNA kit (Macherey-Nagel). cDNA synthesis and qRT-PCR was carried as mentioned above.

### Statistical analysis of data

All the experiments were performed a minimum of three independent times (unless otherwise specified in figure legends) to generate Mean ± SEM that are plotted graphically. For statistical significance, paired end, two-tailed t-test was performed and represented as *P*-values. *, ** and *** indicate *P*-values <0.05>0.01, <0.01>0.001 and <0.001 respectively.

## Supporting information

Complete Supplemental File

## Acknowledgements

Special thanks to Mohan Singh and Hitha G Nair for logistic assistance in the laboratory. Giridharan Govindarajan extended assistance with preparation of polysome graph images. This work was supported by the Department of Biotechnology, Govt. of India (BT/PR3975/MED/29/336/2011). M.J. and D.G. received fellowship from University Grants Commission and Council of Scientific and Industrial Research, both under Govt.of India, respectively.

## Graphical Abstract

HCV infection activates mTORC1. Activated mTORC1 phosphorylates ULK1 that, possibly in association with other factors, suppresses transcription of miR-122. Lower miR-122 levels would in turn affect intracellular HCV RNA abundance. mTORC1 and ULK1 increase abundance of extracellular HCV RNA through unknown mechanisms. Though ULK1 is the prime molecule of autophagy regulation, it does not govern autophagy in Huh7.5 cells.

