## Supplementary material for "mTORC1 Restricts Hepatitis C Virus Replication Through ULK1-mediated Suppression of miR-122 and Facilitates Post-replication Events": Complete Supplemental File

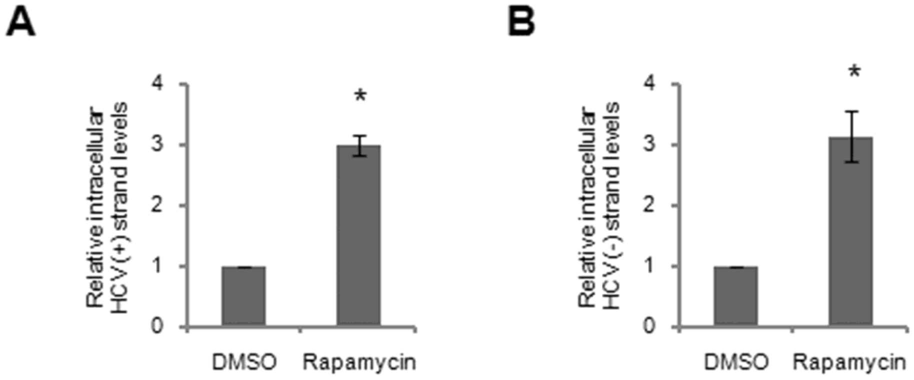


**Figure S1. Increased intracellular HCV RNA abundance upon long-term inhibition with Rapamycin**. Huh7.5 cells were infected with 0.5 MOI HCV and then were treated with 25nM Rapamycin for 24 hrs at 48 hpi respectively. (A) and (B) demonstrate relative intracellular levels of (+) and (-) strands of HCV respectively upon 24 hrs rapamycin treatment, quantified by qRT-PCR.

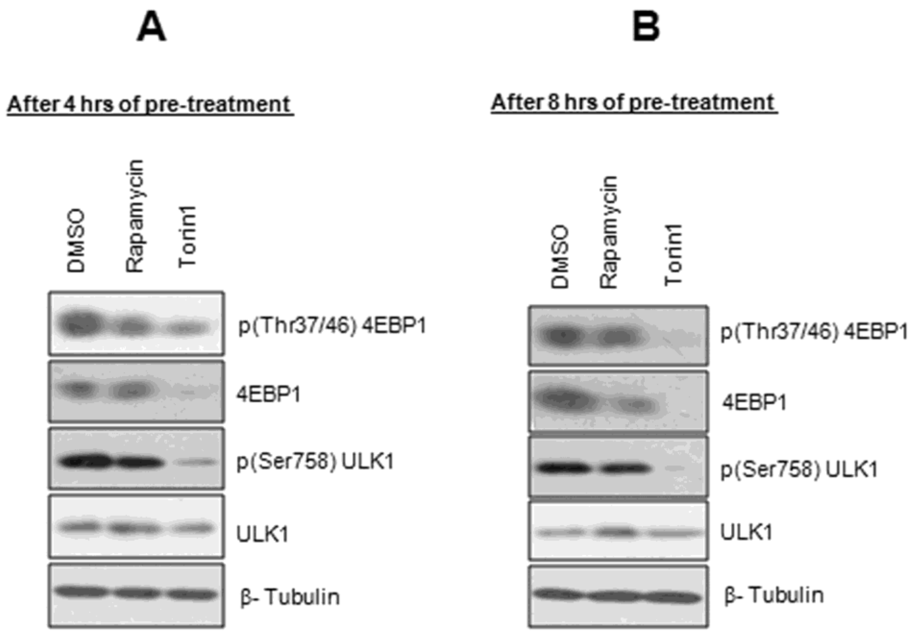


**Figure S2. Inhibition of mTORC1 at the end of 4 hrs and 8 hrs of Torin1 treatment**. (A) Huh7.5 cells were treated with Rapamycin or Torin1 for 4 hrs at the end of which they were harvested for confirmation of mTORC1 inhibition by immunobloting. (B) Similar to (A), but the treatment was extended for another four hours.

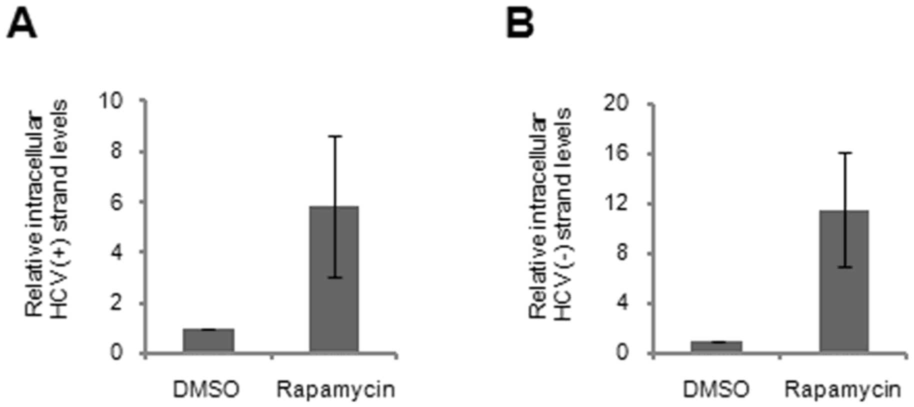


**Figure S3. Effect of pre-inhibition with Rapamycin prior to HCV infection**. As in Fig. 3, cells were pre-treated with Rapamycin 4 hrs before infection. Following this, Rapamycin treatment continued during the 4hrs of HCV infection. Cells were then cultured in fresh media for 68 hrs before harvesting. (A) and (B) depict relative intracellular HCV levels of (+) and (-) strands of HCV respectively.

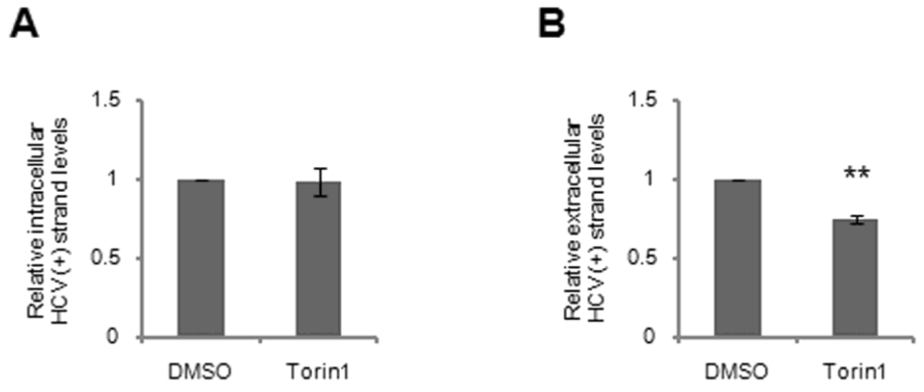


**Figure S4. Effect of mTOR inhibition on HCV post-replication steps.** As in Fig.2, HCV infected cells were inhibited with Torin1 at 48 hpi for 24 hrs, after which the cells were harvested. (A) Relative HCV (+) strand abundance from the secondary infected Huh7.5 cells, infected with intracellular virus of Torin1 treated cells of the primary infection. (B) Relative HCV (+) strand abundance from the secondary infected Huh7.5 cells, infected with the supernatant of Torin1 treated cells of the primary infection.

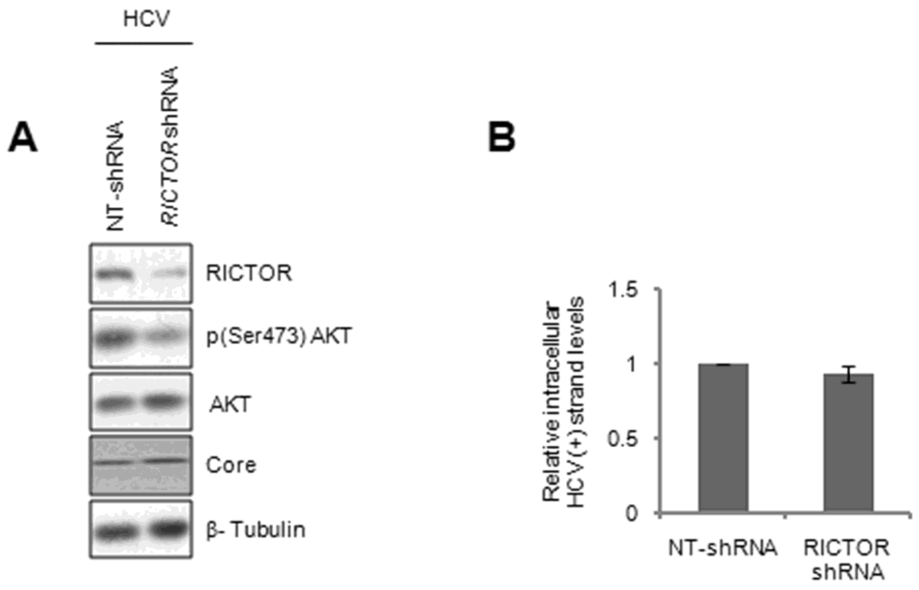


**Figure S5. mTORC2 does not regulate HCV infection.** (A) Huh7.5 cells were transfected with shRNA plasmid pool targeting RICTOR or with non-targeting control shRNAs. 18 hrs later, cells were infected with HCV. Cells were harvested 72 hpi and immunobloted for confirmation. (B) Relative intracellular levels of HCV (+) strands assayed in *Rictor* depleted cells by qRT-PCR.

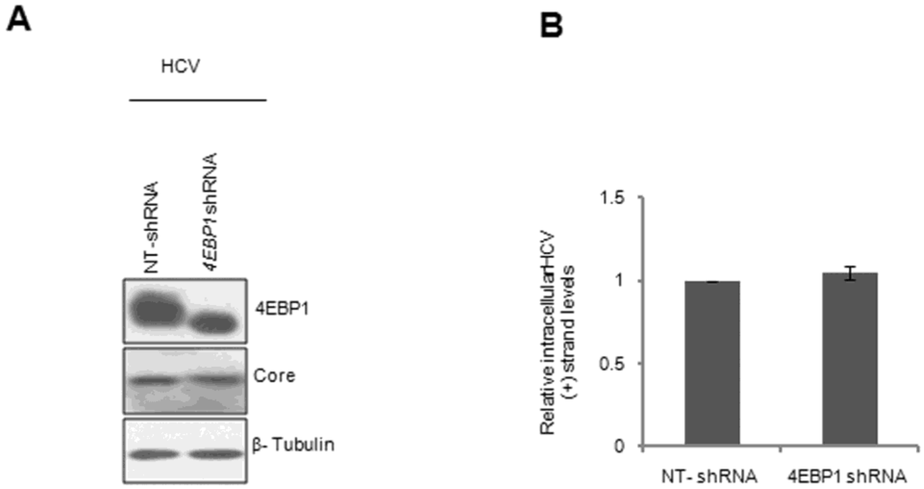


**Figure S6. Role of 4EBP1 in mTORC1 mediated HCV restriction**. (A) Immunoblot demonstrating stable knockdown of 4EBP1 in Huh7.5 cells. Huh7.5 cells were transfected with lentiviral shRNA pool targeting 4EBP1 or with control non-targeting lentivirus pool and were subsequently selected in puromycin. (B) Relative HCV (+) strand levels in 4EBP1-KD Huh7.5 cells as compared with the scrambled control cells.

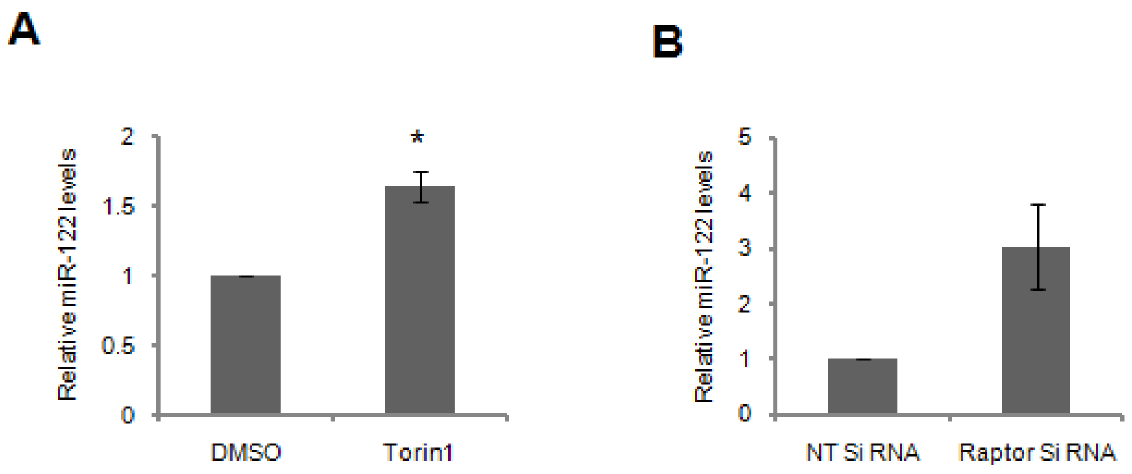


**Figure S7. Effect of Torin1 prolonged-inhibition and *RPTOR silencing* on miR-122 levels**. (A) Huh7.5 cells infected with HCV were treated with Torin1 at 48 hpi for 24 hrs (B) *RPTOR* silencing was performed in Huh7.5 cells followed by HCV infection as explained in Figure 5. The miR-122 levels were quantified by Taqman based kits and normalized against *RNU6B.*

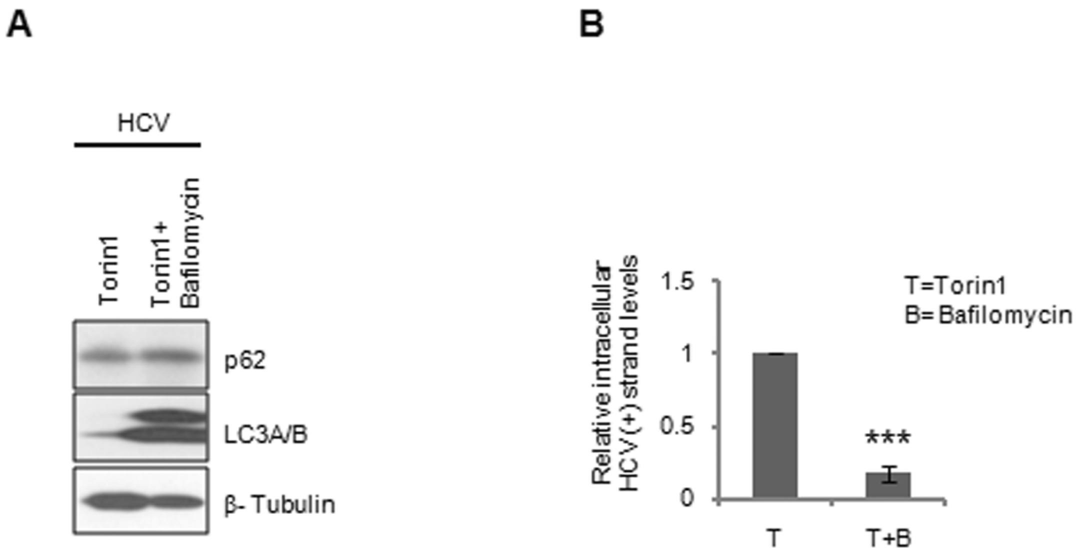


**Figure S8. Bafilomycin overrides effects of Torin1.** (A) Huh7.5 cells were treated with 750nM Torin1 in presence of absence of 50nM Bafilomycin for 6 hrs. Subsequently, the cells were infected with HCV for 4 hrs alongside the inhibitors. 68 hrs later, the cells were harvested and analyzed by immunobloting. (B) represents the relative intracellular HCV (+) strand levels in the treated and the control cells. T =Torin1, B=Bafilomycin.

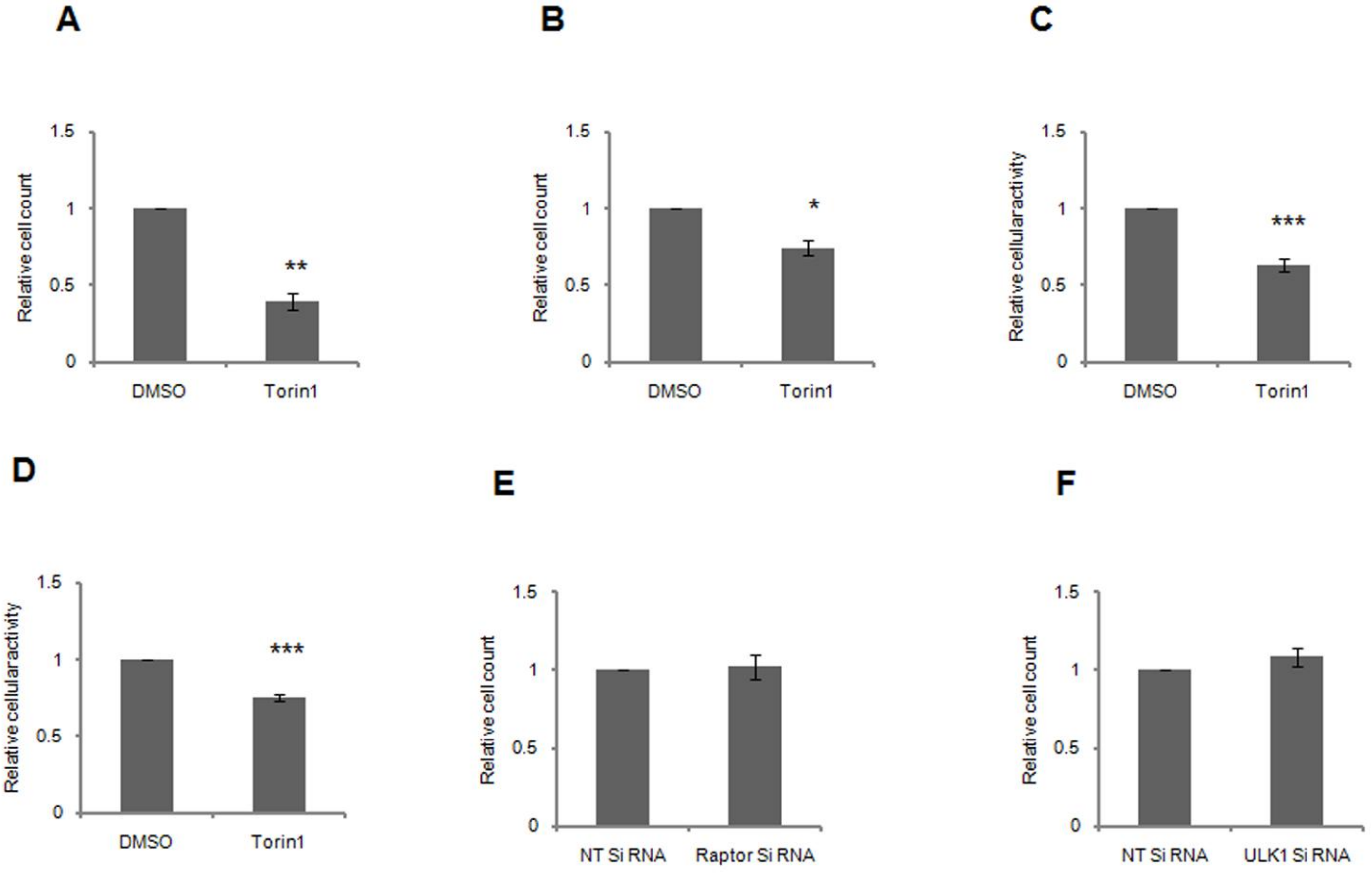


**Figure S9. Relative cell proliferation and viability.** Relative cell proliferation on (A) Preinhibition (B) Prolonged inhibition. Relative cellular activity determined through MTT assay during (C) Preinhibition and (D) Prolonged inhibition Relative cell proliferation on (E) *Rptor* and (F) *ULK1* depletion.

**Table S1. Overview of primers**

| Primer | Description | Sequence |
|---|---|---|
| **HCV RT primer For** | cDNA synthesis primer (- strand) | 5'-ATGAATCACTCCCCTGTGAG-3' |
| **HCV RT primer Rev** | cDNA synthesis primer (+strand) | 5'-TGCACGGTCTACGAGACCTC-3' |
| **HCV SG1For** | Forward primer for RT PCR of HCV | 5'-TATGCCCGGCCATTTGGGCG-3' |
| **HCV SG1Rev** | Reverse primer for RT PCR of HCV | 5'-TACGAGACCTCCCGGGGCAC-3' |
| **GAPDH For** | Forward primer for RT PCR of *GAPDH* | 5′-ATGGGGAAGGTGAAGGTCG-3′ |
| **GAPDH Rev** | Reverse primer for RT PCR of *GAPDH* | 5′-GGGGTCATTGATGGCAACAATA-3′ |
